# Template plasmid integration in germline genome-edited cattle

**DOI:** 10.1101/715482

**Authors:** Alexis L. Norris, Stella S. Lee, Kevin J. Greenlees, Daniel A. Tadesse, Mayumi F. Miller, Heather Lombardi

**Affiliations:** Center for Veterinary Medicine, Food and Drug Administration, Rockville, MD 20855

## Abstract

We analyzed publicly available whole genome sequencing data from cattle which were germline genome-edited to introduce polledness. Our analysis discovered the unintended heterozygous integration of the plasmid and a second copy of the repair template sequence, at the target site. Our finding underscores the importance of employing screening methods suited to reliably detect the unintended integration of plasmids and multiple template copies.

## Article

As genome editing technology evolves, so does our understanding of the unintended alterations it produces, both in form and frequency. Several sequencing-based methods have been developed to screen for off-target errors (GUIDE-Seq^1^, SITE-Seq^2^, CIRCLE-Seq^3^, DISCOVER-Seq^4^), and long-read sequencing of the target site can be used to detect on-target errors. Each screening approach carries assumptions and biases that may allow alterations of unexpected types to go undetected. Recent examples of previously unexpected alterations are complex genomic rearrangements at or near the target site in mammalian genome editing experiments^5, 6^. The complex rearrangements included insertions, deletions, inversions, and translocations that were difficult to detect by standard PCR and DNA sequencing methods.

In this study, we analyzed the target site of publicly-available whole genome sequencing data^7^ from genome-edited calves to confirm the intended edit and to screen for potentially undetected on-target errors (**Supplementary Methods**). The calves were genome-edited^7^, using transcription activator-like effector nucleases (TALENs) and a repair template for homology-directed repair (HDR)^8^, to introduce the Celtic polled allele (*P*_*c*_), a variant that produces the hornlessness (polled) trait in cattle. The *P*_*c*_ variant, common in some cattle breeds^9^, is a 212-bp duplication in place of a 10-bp sequence in an intergenic region on chromosome 1 (chr1:2,429,000-2,429,500; bosTau9). The variant follows an autosomal dominant inheritance, but the mechanism underlying the association with polled trait is unknown.

Given that the repair template plasmid was delivered in the pCR2.1-TOPO plasmid (**Fig. 1a**), we included the plasmid backbone sequence in our comparison of the sequencing reads with the bovine reference genome. In our analysis, we discovered the presence of the full-length plasmid backbone in both genome-edited calves (**Supplemental Fig. 1**). While one allele contained the intended edit, identical to the naturally-occurring *P*_*c*_ variant (**Fig. 1b**), close inspection revealed integration of the plasmid and a second copy of the repair template sequence at the target site in the other allele in both calves (**Fig. 1c**). The plasmid-containing allele (denoted *P*_*c*_*) was found to have been inserted continuous with the template ends, producing a duplication of the template and two novel bovine-plasmid junctions (**Supplemental Fig. 2**). No off-target insertions of the plasmid or the template were detected.

**Fig. 1:**
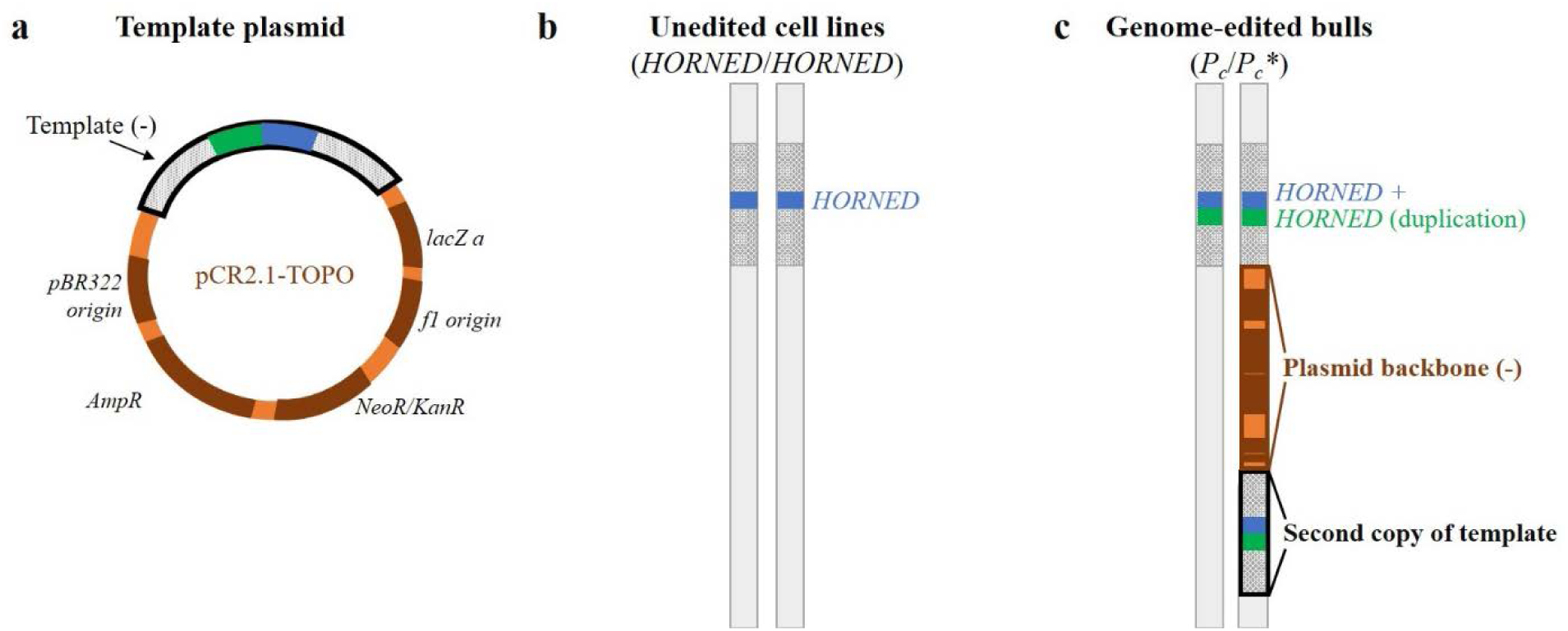
Template plasmid integration at the target site of genome-edited calves. Genomic structure of the template plasmid (**a**), unedited parental cell lines (**b**), and the genome-edited calves (**c**). (**a**) The repair template, containing the *P*_*c*_ sequence and flanking homology arms, is inserted in the pCR2.1 plasmid in an antisense orientation at the TOPO cloning site. (**b**) The unedited parental cell lines are homozygous for *HORNED*. (**a**) The genome-edited calves are heterozygous: one chromosome contains the intended edit (*P*_*c*_), while the other chromosome harbors template plasmid integration, in addition to the intended edit.

Previously, the template plasmid integration was not detected^7^. Probable reasons include: the plasmid backbone was not included in the sequence alignment, elevated noise at the target locus, limited signal of the sequencing data, and PCR conditions insensitive to detect the integrations. The noise was elevated due to the complex sequence context that obscured the integration: (1) the *P*_*c*_ variant itself is a duplication of the reference sequence (*HORNED* allele) in place of a 10-bp sequence, and (2) the target locus is highly repetitive, potentially masking rearrangements. The signal is limited by the sequencing depth of 20 reads for each DNA base, on average. In an ideal scenario, heterozygosity would result in 10 reads identifying the plasmid insertion but given that the plasmid DNA sequence is not in the reference genome, the plasmid reads would remain unmapped. The template plasmid integration was not detected by PCR genotyping^7^ due to the following: (1) the expected PCR amplicons, correctly sized for the *P*_*c*_ variant (212-bp duplication in place of a 10-bp sequence), were produced, (2) the primers were not designed to amplify the plasmid, (3) the amplicons produced by the template plasmid integration were prohibitively large, and (4) the qualitative nature of the assay was insensitive to the increased number of template copies (**Supplementary Methods**, **Supplemental Fig. 3, and Supplemental Table 1**).

Next, we performed a literature search of template plasmid integration at the target site in genome editing experiments, to determine the prevalence of this class of unintended alterations. We found that while there are reports, the template plasmid integration is often not a major finding, and thus we suspect that the integration errors are under reported or overlooked. Template plasmid integration events are known to occur with zinc finger nucleases (ZFNs) at both the target and off-target sites^10, 11^. Using HEK-293 cells, Olsen, *et al.*^10^ showed that transfection of plasmid alone resulted in plasmid integration at a rate of 28×10^−5^ per cell; the addition of one ZFN increased the frequency to 55×10^−5^ per cell and two ZFNs further increased the frequency to 99×10^−5^ per cell. Work by Dickinson *et al.*^12^, using CRISPR/Cas9 with a template plasmid in *C. elegans*, reported the integration of a second copy of the template at the target site. Additional publications using CRISPR/Cas9 with double stranded DNA (dsDNA) repair templates showed that the dsDNA templates can form multimers that integrate into the target site in fish^13^ and mice^14^.

Our discovery highlights a potential blind spot in standard genome editing screening methods. In light of our finding, we propose modifications to current screening methods to enable detection of plasmid integration and integration of multiple template copies. The alignment of sequencing data should include both the reference genome and plasmid sequences. PCR genotyping should incorporate plasmid-specific primers. Methods to detect increased copies of the template and unintended integration of the template plasmid include long-range PCR conditions, quantitative PCR (e.g., digital droplet PCR), Southern blot, and long-read sequencing (e.g., Nanopore, PacBio). The application of suitable screening methods will provide a more precise measure of the prevalence of template plasmid integration events and drive improvements to genome editing, to the benefit of the field.

## Acknowledgments

We acknowledge Recombinetics, Inc. for generating the animals and sharing the sequencing data through NCBI SRA. We appreciate Drs. Alison Van Eenennaam, C. Titus Brown, and Tamer Mansour for helpful discussions concerning our discovery of the template plasmid integration. We thank Drs. Mike Mikhailov and Fu-Jyh Luo for technical expertise, as well as Drs. Harlan Howard, Andrew Fidler, and Kelly Underwood for veterinary expertise.

