## Supplemental Materials for "Template plasmid integration in germline genome-edited cattle"

### Supplementary Methods

*Generation and characterization of the genome-edited calves:*

Carlson *et al.*[1] generated the calves using transcription activator-like effector nucleases (TALENs) mediated genome editing procedure previously performed *in vitro* by Tan *et al.*[2]. For each calf, an unedited fibroblast cell line was transfected with Fok1 nuclease mRNA and a repair template plasmid. The template plasmid was constructed through insertion of the $P_c$-containing DNA template (1,594-bp) in pCR2.1 plasmid (Life Technologies; Carlsbad, CA) at the TOPO insertion site (294-bp position of plasmid). The template was in an inverted orientation to the plasmid (**Fig. 1b**). The edited cell lines were cloned using somatic cell nuclear transfer and the resulting offspring that were then genotypically and phenotypically characterized[1]. Homozygous integration of the $P_c$ allele was confirmed by PCR genotyping (**Supplemental Fig. 3**). The polled phenotype was confirmed by the absence of horned buds by palpation.

*Sequencing data:*

Carlson *et al.*[1] performed whole genome sequencing on the genome-edited calves and their unedited parental cell lines. The four samples were subjected to paired-end (125-bp x 2) sequencing to a 20X average depth of coverage on the HiSeq2500 instrument (Illumina, San Diego, CA). The raw sequencing data was submitted to NCBI's Sequencing Read Archive (SRA Study ID: SRP072240). PCR genotyping confirmed $P_c$ homozygosity of the genome-edited calves. Palpation for the absence of horn buds confirmed the polled trait of the genome-edited calves.

*Sequencing analysis:*

We performed an independent analysis of the raw sequencing data from the two germline genome-edited calves (SRR3290535 and SRR3290615) and their unedited parental cell lines (SRR3290631 and SRR3290632). The sequencing reads were aligned to both the expected locus sequence (reference genome; bosTau9/ARS-UCD1.2) and exogenous DNA delivered for genome editing (pCR2.1-TOPO plasmid), producing a “bosTau9+pCR2.1-TOPO” reference sequence. We included the full bovine reference genome, rather than restricting our analysis to the polled locus, to enable detection of rearrangements involving other regions of the genome and to avoid erroneous mapping of repetitive regions. The sequencing reads were first trimmed (Trimmomatic[3] version 0.38; parameters: LEADING:3 TRAILING:3 SLIDINGWINDOW:4:15 MINLEN:36) and then aligned to bosTau9+pCR2.1-TOPO (BWA-MEM[4] version 0.7.16a; parameters: -M -R "@RG\tID:id\tSM:sample\tLB:lib”). The sequencing reads alignments were then visually inspected using IGV[5] (version 2.4.14).

*Location of Carlson, et al.[1] primers* ***(Supplemental Table 1****) in the repair template (bt9 chr1:2,428,499 -2,429,891):*

atcgaacctgggtcttctgcattggctggcagattctttaccactgagccaccacaccctagagtgcaaaggggggct
tagcaccacaggaggttcacaaagatgtaagctgttcttattaaggctgaggtgggggttgggagaagggggagaaa
aagttttgtaagttgttaatttataataaatccccaaagaaatggtctttcaagtacatacttatctaaaactttgt
caataggggaaatgttcttaggagagaaaaggaatttttttcttttagcataaagctgactttctaatatgggcttcc
ctagtagctcagctggtgaagaacccgcctgcaatgtgggaaacctgggtttgagccctgggttgggaagatcccct
ggagaaaagaatggctacccactccagtattctggcctggagaattccatggactgtataccatggggttgcaaaga
gttggacacgactgagtgactttcattttcactttctgactttctaatattcgaggaatgcttagaagtgtggccgg
tagaaaatagtcctttgtgcctgggacttcaagaaggcggcactatcttgatggaactcagtctcatcacctgtgaa
atgaagagtacgtggtaccaactactttctgagctcacgcacagctggacgtctgcgcctttcttgttatactgcag
atgaaaacattttatcagatgtttgcctaagtatggattacatttaagatacatatttttctttcttgtctgaaagt
ctttgtagtgagagcaggctggaattatgtctggggtgagatagttttcttggtaggctgtgaaatgaagagtacgt
ggtaccaactactttctgagctcacgcacagctggacgtctgcgcctttcttgttatactgcagatgaaaacatttt
atcagatgtttgcctaagtatggattacatttaagatacatatttttctttcttgtctgaaagtctttgtagtgaga
gcaggctggaattatgtctggggtgagatagttttctttgctctttagatcaaaactctcttttcatttttaagtct
atcccaaaagtgtgggaggtgtccttgatgttgaattataggcagagggtcagtttatcaacacccaagaccaacat
ctctgcctttgataagagatagaaatagaagtggagagagaggaggaaaaacatgactcacgatacattctgggttg

60 tttgtttttgtttttatttttgttttggaaggagcgggtgggggaacgtgctgattaaagaaagtctagagagaaca

61 agattcttaaaaataaatccacagtgaagcccagcgggtggggatttcccacagattttcagggctttttttgtgttg

62 tcatggggatattagtcaatgttggtgtcttattttggagtcactatgagtgaaccatgtttaaggagctatggctc

63 agctgctaaactattttcaaaaggaaaatggtgtgttacggtttccgagcagtggggccctggtacaggtaatatca

64 tactcaaaagcactcttttgtgctacatgaa**tgaatcctgctaaaccatgcgga**

**Supplementary Table 1: Primers used by Carlson, *et al.*[1] for PCR genotyping**

| **Primer set** | **Forward primer** (5’-3’) | **Reverse primer** (5’-3’) | ***HORNED* amplicon** (bp) | **$P_c$ amplicon** (bp) |
|---|---|---|---|---|
| **btHP** (overlap) | GAAGGCGGCACTATCTTGATGGAA | GGCAGAGATGTTGGTCTTGGGTGT | 389 | 591 |
| **HP1748** (flank) | GGGCAAGTTGCTCAGCTGTTTTTG | TCCGCATGGTTTAGCAGGATTCA | 1,546 | 1,748 |

**Colored for their position in the repair template sequence (**Supplemental Methods**)*

### Supplemental Figures

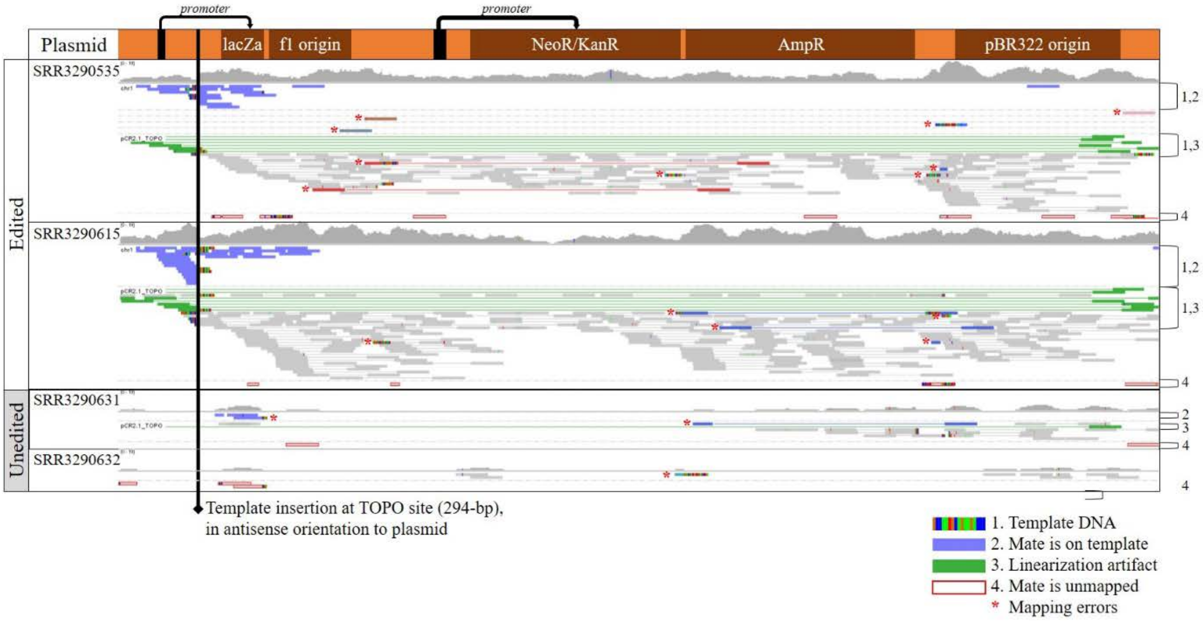


**Supplemental Fig. 1: Plasmid DNA is present in genome-edited calves.**

Shown are all sequencing reads that aligned to the linearized plasmid backbone, with origins, genes, and gene promoters listed. For each sample, the coverage track is shown above the individual sequencing read pairs, for which lines connect the mate pairs. The template-plasmid junctions are indicated by discordantly mapped pairs and split reads. In discordant pairs, the read's mate is aligned to the template, not the plasmid. In split reads, the mismatched bases at an end of the read are the template sequence, because the read spans the template-plasmid junction. In both genome-edited calves, the reads span the entire length of the plasmid, and at a coverage consistent with heterozygosity (~10X coverage). The unedited parental cell lines have sparse, low-level coverage of the plasmid, consistent with contamination. The coverage track scale is uniform across samples (0-19 reads). Reads at the start and end of the plasmid sequence are artifacts of linearization of the circular plasmid.

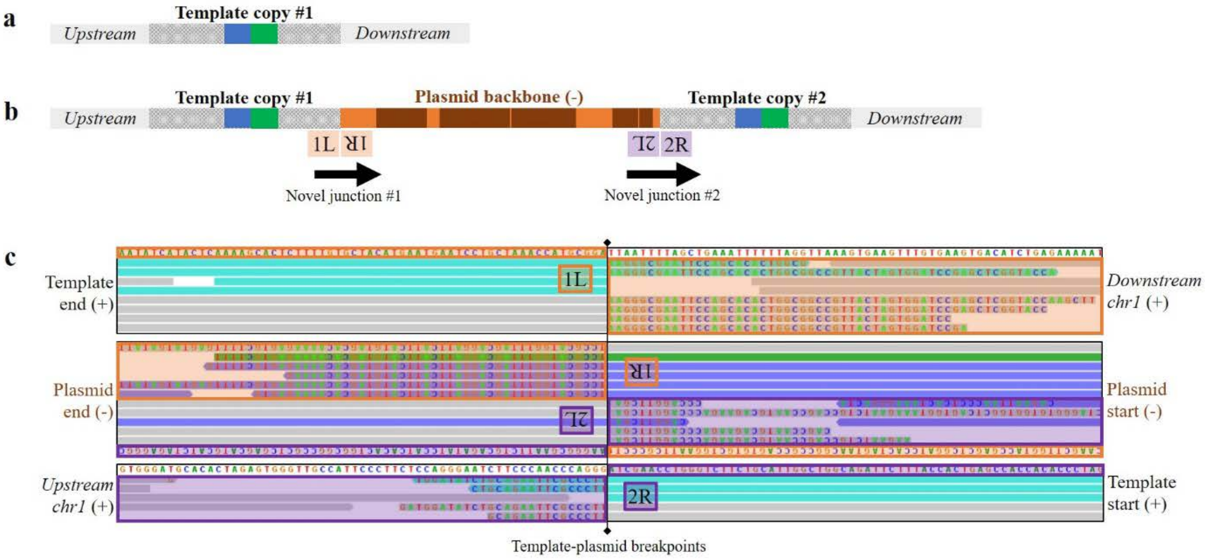


**Supplemental Fig. 2: Template plasmid is integrated at the target site in genome-edited calves.**

Compared to the naturally occurring $P_c$ sequence (**a**), the chromosome with the template plasmid integration (**b**) adds two novel junctions created by the plasmid and flanking template sequences. The junctions are evidenced by the aligned sequencing reads, shown here for a representative sample (SRR3290615) in IGV (**c**). The plasmid sequence is inverted as a result of the antisense insertion of the repair template at the TOPO site during cloning. For each template-plasmid junction, the left (L) and right (R) breakpoints are indicated. The first novel junction's breakpoints are the end of the first template copy (bt9 chr1:2,429,911; 1L) with the start of the plasmid (pCR2.1-TOPO:294-bp(-); 1R). The second novel junction's breakpoints are the end of the plasmid (pCR2.1-TOPO:295-bp(-); 2L) and the start of the second template copy (bt9 chr1:2,428,500; 2R).

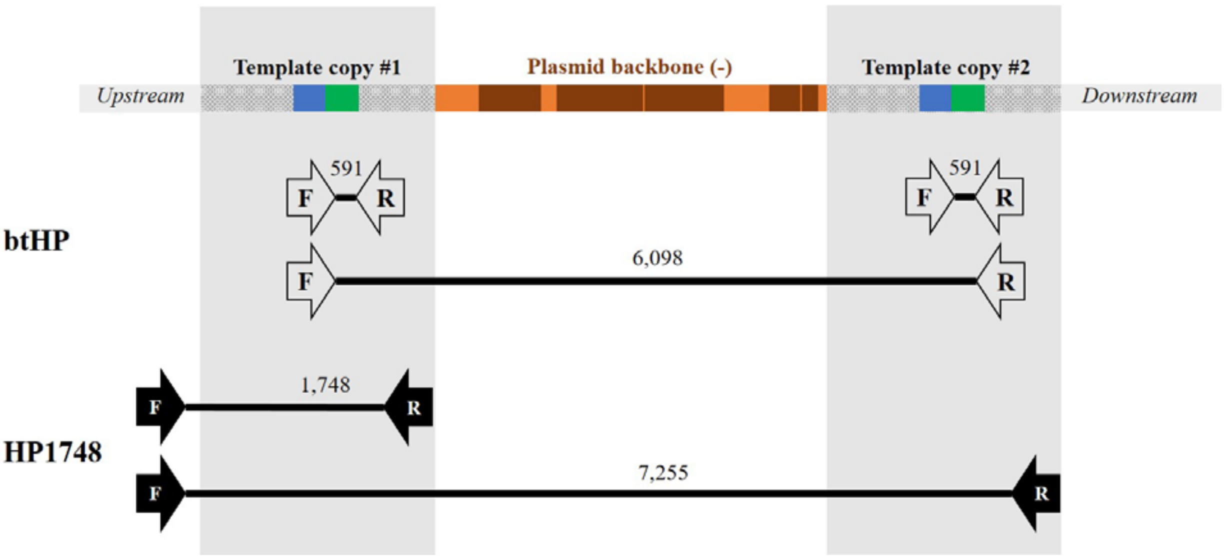


**Supplemental Fig. 3: Template plasmid integration was not detected by PCR genotyping.**

Carlson, *et al.*[1] used two primer sets, btHP and HP1748 (**Supplemental Table 1 and Supplemental Methods**), for confirming $P_c$ homozygosity of the genome-edited calves. Neither primer set detected the template plasmid integration by gel electrophoresis or Sanger sequencing of the TOPO-cloned amplicon. The btHP primers overlapped $P_c$, while the HP1748 primers flanked $P_c$. In HP1748, the forward primer was 154-bp upstream of the start of the repair template, and the reverse primer at the end of the repair template. For both primer sets, amplicons containing the plasmid would be prohibitively large (6,098-bp and 7,255-bp, respectively). With regards to the second copy of the repair template, the doubled amount of $P_c$ amplicons for primer set btHP (591-bp) would be difficult to detect without a quantitative method, and the HP1748 primer set does not amplify the second copy because the HP1748 forward primer was upstream of the repair template.
